# Uncovering hidden gene-trait patterns through biclustering analysis of the UK Biobank

**DOI:** 10.1101/2024.11.08.622657

**Authors:** Milton Pividori, Suraju Sadeeq, Arjun Krishnan, Barbara E. Stranger, Christopher R. Gignoux

**Author notes:** Correspondence possible via GitHub Issues or email to Milton Pividori < >. Funded by The National Human Genome Research Institute (K99/R00 HG011898); The Eunice Kennedy Shriver National Institute of Child Health and Human Development (R01 HD109765). @. compbiologist. popgenepi.

## Abstract

The growing availability of genome-wide association studies (GWAS) and large-scale biobanks provides an unprecedented opportunity to explore the genetic basis of complex traits and diseases. However, with this vast amount of data comes the challenge of interpreting numerous associations across thousands of traits, especially given the high polygenicity and pleiotropy underlying complex phenotypes. Traditional clustering methods, which identify global patterns in data, lack the resolution to capture overlapping associations relevant to subsets of traits or genes. Consequently, there is a critical need for innovative analytic approaches capable of revealing local, biologically meaningful patterns that could advance our understanding of trait comorbidities and gene-trait interactions. Here, we applied BiBit, a biclustering algorithm, to transcriptome-wide association study (TWAS) results from PhenomeXcan, a large resource of gene-trait associations derived from the UK Biobank. BiBit allows simultaneous grouping of traits and genes, identifying biclusters that represent local, overlapping associations. Our analyses uncovered biologically interpretable patterns, including asthma-related biclusters enriched for immune-related gene sets, connections between eye traits and blood pressure, and associations between dietary traits, high cholesterol, and specific loci on chromosome 19. These biclusters highlight gene-trait relationships and patterns of trait co-occurrence that may otherwise be obscured by traditional methods. Our findings demonstrate that biclustering can provide a nuanced view of the genetic architecture of complex traits, offering insights into pleiotropy and disease mechanisms. By enabling the exploration of complex, overlapping patterns within biobank-scale datasets, this approach provides a valuable framework for advancing research on genetic associations, comorbidities, and polygenic traits.

## Introduction

The availability of genome-wide association studies (GWAS) and large-scale biobanks, which house extensive genomic and phenotypic data, has revolutionized our ability to investigate the biology underlying complex traits and diseases [1,2]. These resources allow researchers to identify genetic loci associated with various traits, offering a broad view of genetic influences on disease susceptibility and comorbidity. However, the immense scale of GWAS data from diverse populations and phenotypes presents challenges for interpretation and synthesis [3]. As more GWAS findings accumulate, understanding how these genetic variants contribute to the polygenic and pleiotropic nature of complex traits becomes increasingly critical [4]. This complexity necessitates innovative analytical approaches to uncover and interpret subtle, biologically meaningful relationships within large datasets.

Traditional clustering methods, such as *k*-means or hierarchical clustering, have been widely applied to biological data to group traits or genes based on global patterns [5,6,7]. These methods have been effective in identifying broad patterns across entire datasets, grouping either samples or genes according to shared characteristics. However, complex traits and diseases often exhibit polygenic architectures where specific genes interact with multiple traits, and certain traits share overlapping genetic influences. Traditional clustering fails to capture these local, overlapping patterns [8], especially in high-dimensional data, where subsets of genes might only be relevant to subsets of traits. In such cases, biclustering approaches [9] offer a solution by simultaneously grouping genes and traits, identifying local patterns that reflect biological specificity. While some studies have demonstrated the effectiveness of biclustering in gene expression analysis [10,11], its application to GWAS or TWAS data remains limited, leaving an unmet need for methods that can disentangle the intricate connections within these associations.

In the realm of complex traits, transcriptome-wide association studies (TWAS) offer an additional layer of insight by linking genetic variants to gene expression and, subsequently, to traits [12,13,14,15]. TWAS integrates gene-trait associations across tissues, enhancing our ability to infer potential causal genes and pathways underlying GWAS loci. However, even with TWAS, it remains challenging to interpret the vast array of gene-trait associations in a biologically meaningful way. Although TWAS has been instrumental in identifying genes involved in individual traits [16], understanding how these genes contribute to trait comorbidity and pleiotropy across many traits and tissues requires an advanced analytical approach that can manage high-dimensional, interdependent data relationships. Therefore, a key gap in the field is the need for methods that can capture these overlapping, complex associations in a way that supports the biological interpretation of TWAS results, particularly across the wide phenotypic spectrum represented in biobank-scale datasets.

In this study, we address this gap by applying a biclustering approach to TWAS results from PhenomeXcan [17], an extensive resource derived from the UK Biobank [2]. Our method uses BiBit [18], a biclustering algorithm that simultaneously groups traits and genes based on local patterns of association. This approach enables the detection of overlapping biclusters where a subset of genes may be linked to multiple traits and vice versa, offering a nuanced view of gene-trait interactions. Our analyses reveal several biologically relevant biclusters, such as those connecting immune-related gene sets to asthma diagnoses, linking eye traits with blood pressure, and associating dietary and cholesterol traits with genes on chromosome 19. These findings demonstrate the utility of biclustering for uncovering patterns that enhance our understanding of complex trait architecture and open new avenues for exploring disease mechanisms and comorbidities.

## Results

### Overview of the biclustering approach

To explore patterns within TWAS results from the UK Biobank, we utilized PhenomeXcan [17], a comprehensive resource of gene-trait associations based on 4,091 GWAS summary statistics from publicly available data [19] and the UK Biobank [2]. We applied the S-MultiXcan approach [14], a cross-tissue method that aggregates gene-trait associations across tissues from PrediXcan [12,13], enhancing statistical power for gene-trait associations. This yielded an association matrix containing *p* -values for 4,091 traits and 22,515 genes.

To capture local patterns of gene-trait associations, we applied the BiBit biclustering algorithm, which detects optimal biclusters in a binarized gene-trait association matrix (see Figure 1a). We binarized the matrix using a Bonferroni-corrected *p*-value threshold (5.49 × 10^−10^), allowing BiBit to detect associations where all gene-trait pairs in each bicluster met this significance threshold. BiBit yielded 20,494 biclusters, with each bicluster representing gene and trait subsets that share a statistically significant association. The resulting data and interactive exploration tools are available at https://pivlab.github.io/biclustering-twas/.

**Figure 1:**
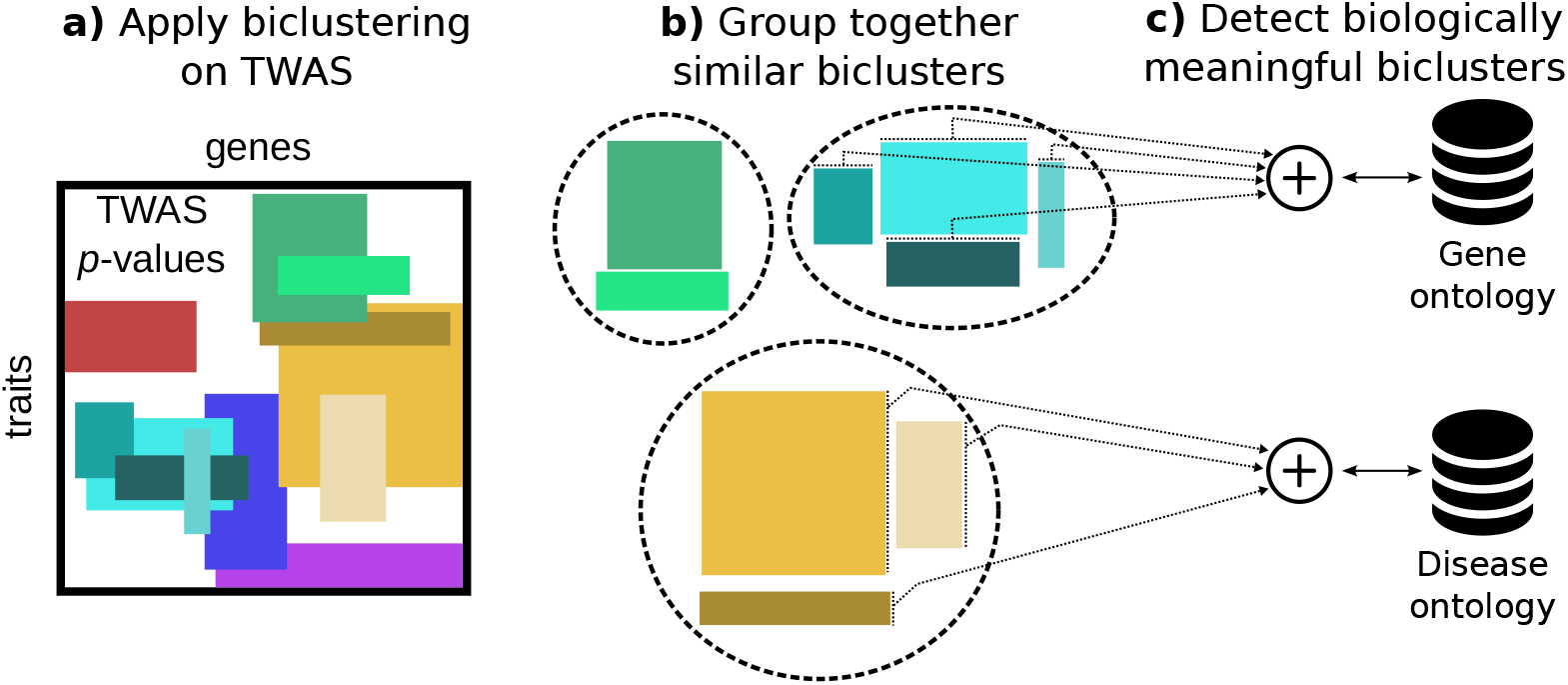
Schematic of the computational analyses. **a)** The BiBit biclustering algorithm was applied to the S-MultiXcan association matrix to identify biclusters—rectangles where subsets of traits (rows) are associated with subsets of genes (columns). **b)** Biclusters were grouped based on gene overlap using a clustering algorithm on the Jaccard similarity coefficient. **c)** Gene Ontology (GO) and Disease Ontology (DO) over-representation analysis was performed on grouped biclusters to assess biological pathway representation.

### Asthma-related biclusters are enriched for immune-related gene sets

To determine if the biclustering approach could capture biologically meaningful patterns, we focused on biclusters containing asthma-related traits. Given the overlap in biclusters, we grouped similar biclusters into meta-biclusters based on gene overlap (Figure 1b). Our steps included: 1) selecting biclusters with asthma-related traits (self-reported asthma, ICD-10 codes J45/J46, and age of asthma onset) and at least 10 associated genes, 2) calculating the Jaccard similarity coefficient for pairwise bicluster comparisons, 3) clustering biclusters with high gene overlap, yielding 10 meta-biclusters, and 4) performing over-representation analyses for Gene Ontology (GO) terms (for genes) and Disease Ontology (DO) terms (for traits) to assess distinct biological mechanisms for asthma.

As shown in Figure 2, each meta-bicluster was enriched for unique GO and DO terms. Nine of the ten meta-biclusters demonstrated distinct GO enrichment, suggesting diverse biological pathways contributing to asthma. For example, meta-bicluster C6 included traits such as celiac disease, systemic lupus erythematosus, and type 1 diabetes, with genes localized to the HLA region on chromosome 6, underscoring its role in autoimmunity. Other meta-biclusters, such as C8, were associated with allergic diseases and the age of asthma onset, with strong links to the 17q12-21 locus, a well-established region for early-onset asthma [20,21,22]. Consistent with a large study on pleiotropy [4], these results suggest that a combination of genes seems to be specific to a particular combination of traits.

**Figure 2:**
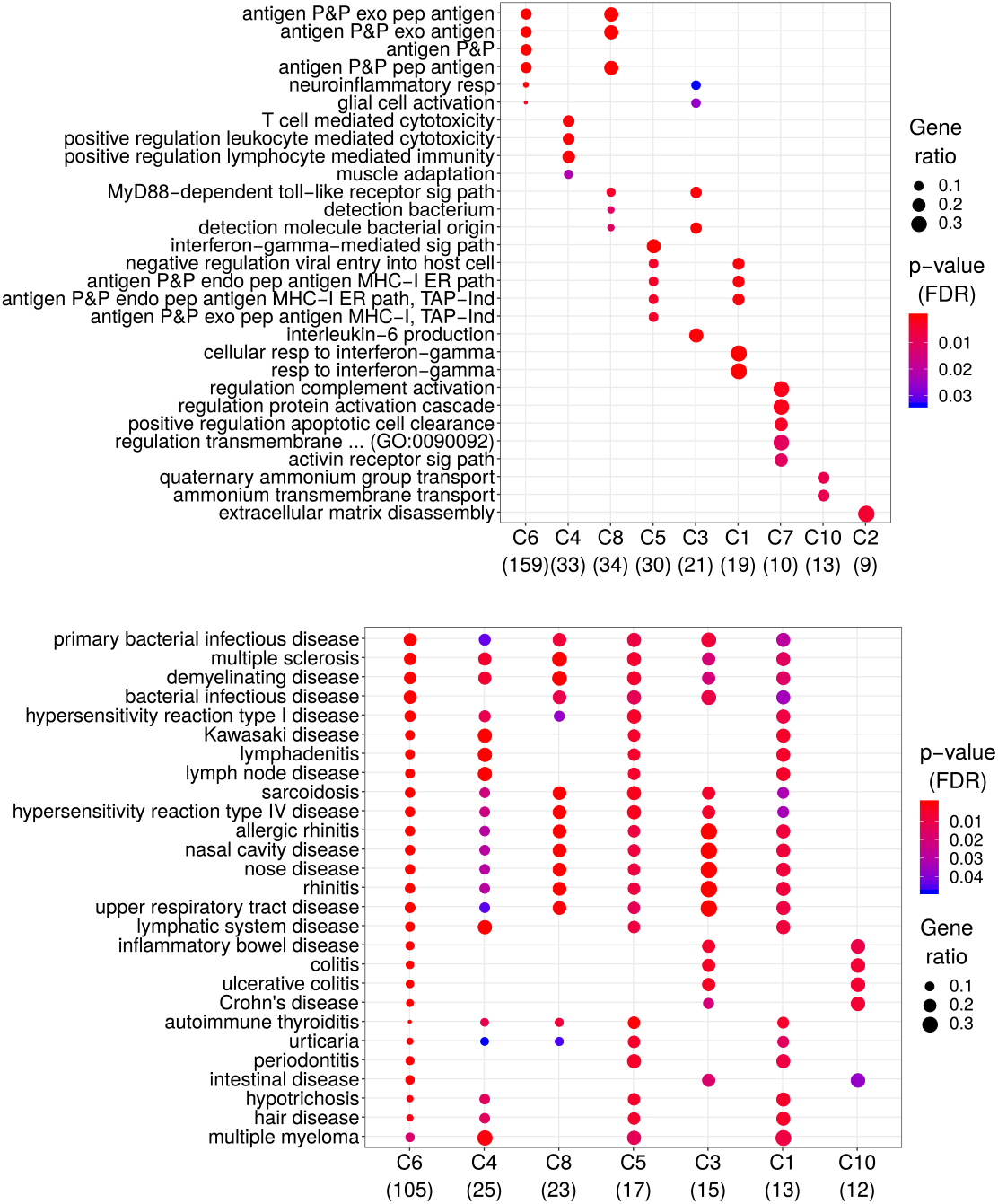
Functional enrichment of asthma-related biclusters. Enrichment analyses of 10 asthma-related meta-biclusters revealed distinct GO terms across genes (top) and DO terms across traits (bottom), illustrating pathway and disease enrichment for asthma susceptibility.

### Biclusters associated with eye measurements

We next investigated biclusters linked to eye-related traits to understand potential gene-trait connections in ocular health. Two notable biclusters involved traits like “Age started wearing glasses” and keratometry measurements alongside blood pressure.

In Figure 3, the left table shows the bicluster associated with “Age started wearing glasses” and keratometry measures, while the right table displays associations with blood pressure. The gene **TXLNB**, involved in muscle function [23], was notable in these associations, aligning with research linking blood pressure with eye health [24]. These findings indicate potential links between eye traits and blood pressure mediated by genes influencing muscular and vascular function.

**Figure 3:**
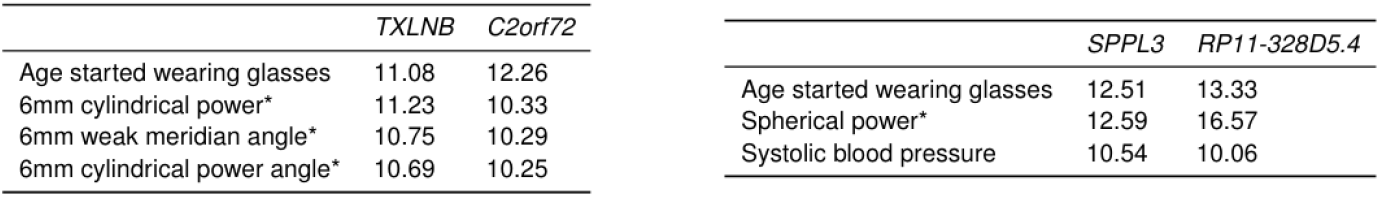
Biclusters associated with eye measurements. Two biclusters showing associations between “Age started wearing glasses” and keratometry measurements (left), and blood pressure (right). Eye traits are marked with an asterisk; cells contain *z*-scores derived from *p*-values.

### Biclusters associated with anthropometric traits

We also identified biclusters related to anthropometric traits, which included a diverse array of body composition metrics.

As shown in Figure 4, this bicluster includes 15 traits related to body composition, including BMI, weight, and measurements of fat and lean mass. Key genes, such as *TFAP2B* and *USP36*, have roles in adipogenesis and metabolic regulation, suggesting involvement in body composition pathways. These genes support the hypothesis that specific genetic networks influence distinct aspects of anthropometric traits, including fat and lean mass distribution.

**Figure 4:**
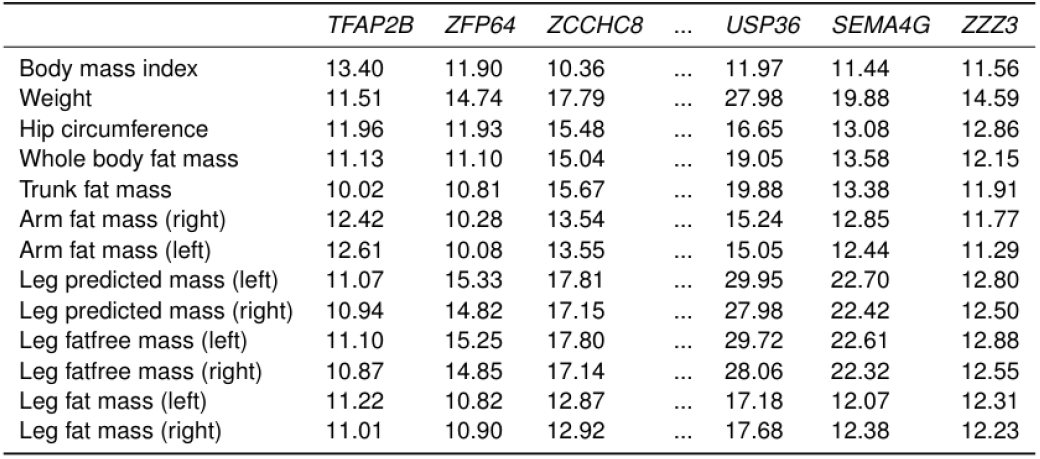
Bicluster associated with anthropometric traits. A bicluster showing associations between 15 anthropometric traits, such as BMI and hip circumference, and 68 genes, a subset of which are shown. Cells contain *z*-scores derived from *p*-values.

### Biclusters associated with high cholesterol and dietary traits

Finally, we identified biclusters connecting high cholesterol with dietary and metabolic traits.

Figure 5 displays a bicluster associated with high cholesterol, cholelithiasis, and dietary traits, including processed meat intake, fish intake, and sodium levels. Genes in this cluster, such as *RASIP1, FUT1, FUT2*, and *IZUMO1*, are located within a gene-dense region on chromosome 19. This region, associated with macronutrient intake through genes like *FGF21* [25], suggests links between diet and chronic conditions such as obesity and diabetes. This bicluster highlights how dietary factors may interact with cholesterol metabolism, offering insights into gene-diet interactions relevant to metabolic health.

**Figure 5:**
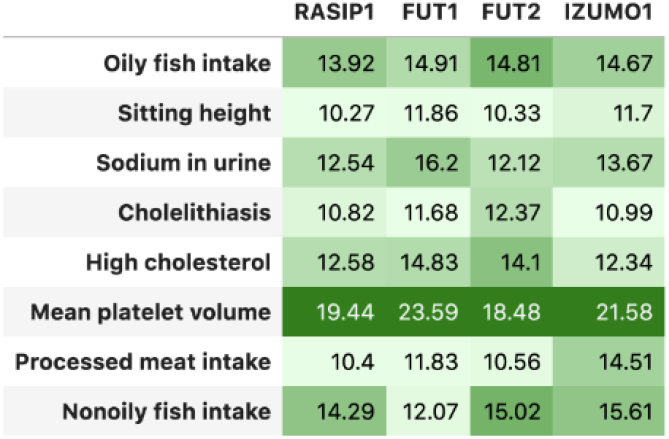
Bicluster associated with high cholesterol and dietary traits. This bicluster shows associations among high cholesterol, cholelithiasis, dietary intake, and specific genes located on chromosome 19. Cells are *z*-scores derived from *p*-values, with conditional formatting to highlight significance.

These results demonstrate that biclustering provides a framework to identify biologically relevant gene-trait relationships, supporting the exploration of complex traits and comorbidities within large biobank datasets.

## Discussion

This study leverages a biclustering approach, specifically the BiBit algorithm, to reveal complex associations within TWAS data, providing insights into the genetic architecture of multiple traits in the UK Biobank. By capturing both gene-trait and trait-trait patterns, we uncovered biologically relevant associations such as asthma-related immune gene clusters, eye trait associations with keratometry and blood pressure, and anthropometric traits connected to genes involved in adipogenesis and metabolic regulation. These findings underscore the potential of biclustering to enhance our understanding of pleiotropy and trait comorbidities, offering a novel framework for exploring the polygenic and interconnected nature of complex traits.

One strength of our study is the use of biclustering to detect local patterns that traditional clustering methods may overlook. While traditional clustering captures only global patterns, biclustering allows for the simultaneous grouping of genes and traits, revealing gene sets associated with specific trait subgroups. This flexibility is particularly beneficial in complex, polygenic traits like asthma, where subsets of genes and phenotypes contribute to distinct pathways. While this approach successfully identified asthma-related clusters enriched for immune-related pathways, the reliance on binary input and a Bonferroni-corrected *p*-value threshold may limit the detection of more subtle associations.

Nonetheless, the identified asthma biclusters are consistent with previous studies on autoimmune and allergic pathways, supporting the validity of our findings.

A limitation to consider is that our biclustering approach does not incorporate tissue specificity, which can affect gene-trait associations in TWAS data. The integration of tissue-specific data could refine the resolution of our biclusters and enhance the biological relevance of the findings. For instance, incorporating tissue information could help distinguish genes that influence systemic traits from those affecting tissue-specific phenotypes, such as eye or respiratory conditions. Future studies could address this limitation by using tissue-aware biclustering methods or incorporating multi-omic data to capture more nuanced layers of biological regulation.

Additionally, while we observed biclusters associating anthropometric traits with genes like *TFAP2B* linked to skeletal and adipogenic pathways, our findings rely on existing TWAS data that may reflect genetic correlations rather than causal relationships. Although TWAS improves on GWAS by prioritizing likely causal genes, causal inference remains a challenge due to potential confounders and linkage disequilibrium effects. Future work could employ Mendelian randomization or fine-mapping to validate the causal roles of identified genes within these biclusters, thereby strengthening the biological interpretations of our findings.

Despite these limitations, our study makes a significant contribution to the field by demonstrating how biclustering can uncover multi-dimensional patterns in TWAS data, an approach that is highly adaptable to other biobank-scale datasets. By revealing new connections between genes and phenotypes, this method provides a foundation for future research on genetic correlations, pleiotropy, and disease comorbidity, fostering new avenues for understanding the biological pathways underpinning complex traits. Moreover, the interactive online resource we provide allows other researchers to explore the identified biclusters, potentially catalyzing novel hypotheses and collaborative efforts to validate and expand upon our findings.

In conclusion, this work advances the field by presenting an innovative analytical framework that bridges gaps in current GWAS and TWAS studies, empowering researchers to interpret and synthesize the large volumes of data from biobanks like the UK Biobank. Our approach holds promise for accelerating biomedical discovery by facilitating the identification of complex trait architectures and may serve as a valuable tool for exploring the polygenic basis of traits and diseases in the post-GWAS era.

## Methods

### Gene-trait associations with TWAS

We used PhenomeXcan [17], a large resource of TWAS results with gene-tissue-trait associations across 4,091 diseases and traits from the UK Biobank [2] and other cohorts, 49 tissues from GTEx, and more than 22,000 genes. PhenomeXcan was built using publicly available GWAS summary statistics from Neale’s lab (GWAS round 2 [19]) to compute 1) gene-based associations with the PrediXcan family of methods [12,13,14], including Summary-PrediXcan (S-PrediXcan) [13], which generates gene-tissue-trait associations, and Summary-MultiXcan (S-MultiXcan) [14], which combines tissue-specific results from S-PrediXcan to generate gene-trait associations; and 2) a locus regional colocalization probability (RCP) between GWAS loci and *cis*-eQTL (i.e., the overlapping of prioritized causal variants for a trait and for a gene’s expression) with fastENLOC, a Bayesian hierarchical model for large-scale data [17,26]. The PrediXcan family of methods first builds prediction models using data from the Genotype-Tissue Expression project (GTEx v8) [27] for gene expression imputation and then correlates this predicted expression with the phenotype of interest.

### Biclustering analysis

We used a biclustering method that can jointly model pleiotropic genes and polygenic traits by allowing overlapping biclusters. For this, we applied the BiBit algorithm [18] on gene-trait associations from PhenomeXcan.

Since BiBit works on binary data, given a matrix *M* ^*n*×*m*^ (*n* = 4,091 traits and *m* = 22,515 genes) with *p*-values from S-MultiXcan, we defined the binary matrix *E*^*n*×*m*^, with *E*_*ij*_ = 1 if *M*_*ij*_ < *t*, and *E*_*ij*_ = 0 otherwise, where *t* was set to a Bonferroni significant *p*-value (5.49 × 10^−10^). A bicluster *ℬ*_*ℓ*_ = (𝒯_*ℓ*_, 𝒢_*ℓ*_) corresponds to a subset of traits 𝒯_*ℓ*_ ⊂ {1, …, *n*} that are all jointly associated with a subset of genes 𝒢_*ℓ*_ ⊂ {1, …, *m*}, such that *M*_*ij*_ < *t*, ∀*i* ∈ 𝒯_*ℓ*_, ∀*j* ∈ 𝒢_*ℓ*_.

### Hierarchical clustering of biclusters and functional analyses

Biclusters across the entire set of results can have different degrees of overlap, which means that they can be grouped into several clusters. These clusters can be summarized by the frequency of phenotypes and genes across their biclusters. Thus, we proceeded as follows: 1) select all biclusters containing at least 10 genes and a disease of interest, such as an asthma diagnosis (self-reported asthma, ICD-10 codes J45/J46, and age of asthma onset); 2) perform a pairwise comparison of all these biclusters using the Jaccard similarity coefficient over their associated genes; 3) run a hierarchical clustering algorithm (average linkage) and obtain 10 groups of biclusters; 4) perform an over-representation analysis of Gene Ontology (GO) terms to compare those 10 groups of biclusters and assess whether their genes represent known and distinct biological mechanisms for the disease of interest; the same analysis can be done using the Disease Ontology (DO) terms on the traits.

